# Ontology-driven software engineering using LLMs for knowledge graphs in engineering biology

**DOI:** 10.64898/2026.05.29.728869

**Authors:** Ihsan Tolga Medeni, Metehan Ünal, Roberto Galizi, Bryan Bartley, Jacob Beal, Chris J. Myers, Prashant Vaidyanathan, Göksel Mısırlı

## Abstract

Large language models have transformed software engineering practices. However, generated artefacts are not always developer-friendly and may partially meet complex requirements. As the need to standardise, integrate, and develop tools in engineering biology increases, novel approaches are needed to create and maintain intuitive software sustainably. Here, we present an ontology-driven approach using large language models to create user-facing software libraries for knowledge graphs. We introduce an ontology-to-language framework to systematically map domain terms and graph structures. We then demonstrate this approach by creating an ontology for the latest Synthetic Biology Open Language standard and generating the sbol-script software library, which can be used within browsers or to develop applications with native web support. This ontology-driven software engineering approach and these resources are essential for the community and to facilitate the development of sustainable software projects. The SBOL3 Ontology and the sbol-script library are available from https://github.com/SynBioDex/sbol-owl3 and https://github.com/SynBioDex/sbol-script.

## 1 Introduction

Software automation is an important feature for large-scale and self-evolving information systems. Model-driven software engineering (MDSE) can be used to automate the creation of software artefacts from well-defined rules, in which models describe domain-specific requirements (*1, 2*). With the advances in artificial intelligence (AI), there are significant opportunities to bridge MDSE and large language models (LLMs). However, LLMs can generate outputs regardless of the desired outcome and efficiency of the resulting artefacts. AI inconsistency, incompleteness and hallucination are ongoing issues (*3, 4*). There is a need to formally describe requirements for better prompts with well-defined inputs, especially in domains such as engineering biology, where new tools are actively being developed.

Engineering biology workflows may involve several tools, for example, to create, visualise, store, and model biological designs (*5*). Hence, standardisation is essential to enable interoperability between these tools. The Synthetic Biology Open Language (SBOL) has been developed as a community standard to represent biological designs unambiguously and facilitate data exchange across various tools, repositories and workflows (*6, 7*). It can be used to describe biological parts, their interactions, hierarchical designs, and associated sequences and metadata. As engineering biology workflows become increasingly data focused and distributed, SBOL has an important role to support reusability and integration (*8*). SBOL3 is the latest version of the standard (*7*) and builds on graph-based serialisation using the Resource Description Framework (RDF) (*9*). This approach facilitates integration with existing biological knowledge graphs and ontologies. Moreover, it enables the use of readily available semantic web tools and methods to handle, store, and query data (*10*).

Despite these technological foundations, software support for SBOL3 is limited. It is essential to create software libraries that provide an abstraction between tool developers and the detailed SBOL specifications. Existing implementations, such as libSBOLj3 (*11*) and pySBOL3 (*12*), demonstrate the importance of such libraries for wider adoption but also highlight the challenges of creating and maintaining domain-specific software infrastructures. Additionally, developers may prefer different programming languages, such as TypeScript or JavaScript, especially for native web development. Creating new libraries and maintaining them through incremental specification changes for different programming languages clearly benefits from automation.

LLMs are attractive to provide model-to-software coverage flexibly in order to translate heterogeneous knowledge sources into implementable software artefacts with significantly less manual effort. LLMs can also simultaneously process formal descriptions, additional documentation, and use cases through iterative feedback. As a result, LLMs are suitable especially for domains where requirements are semi-structured. This flexibility is important for developing software in engineering biology, where the goal is not only to reproduce syntax, but also to capture relationships, constraints and abstractions that focus on user-facing and maintenance requirements.

Despite their advantages, LLMs can be hallucinatory and produce inconsistent results. Moreover, prompt-oriented approaches are insufficient to fully capture the semantics for com-plex scenarios. These issues require complementing LLMs with a formal representation layer to guide the output generation process. Specification-driven code generation has already been recognised (*13* –*16*) as an approach to bridge the gap between AI systems and actual requirements. Ontology-driven approaches are also considered in LLM applications, as ontologies are machine processable and interpretable (*17* –*19*). However, existing approaches typically use ontologies to train LLMs or link domain entities. The creation of reusable libraries for software developers requires further consideration and should be supported via additional transformation layers to take advantage of the explicit link between syntax and semantics in graph-based models.

Ontologies are ideal to provide explicit and machine-accessible definitions of terms and their relationships for a domain of interest (*20*). Unlike traditional data models, ontologies provide generalised and extensible solutions to model various types of constraints using the rich expressivity of the Web Ontology Language (OWL) and the features of description logics (*21, 22*). Ontological models were also defined for the previous SBOL2, demonstrating the potential of such a semantic layer (*23, 24*). As the biological ontologies are widely used, they can guide the creation of software libraries efficiently.

Ontologies have already been utilised in MDSE even before LLMs to create graph-based libraries (*25* –*27*). In this approach, the process of creating software artefacts can be viewed as data model transformations, mediated by metamodel descriptions from ontologies (*2, 28, 29*). In a typical MDSE workflow, data schemas are converted into software building blocks, such as classes, and the resulting entities are annotated with bindings that can be used to instantiate objects (*30*). Similarly, graph-based approaches (*31*) can use ontology-driven RDF bindings to control how classes and properties are serialised as a set of nodes and edges. Although such frameworks (*31*) were previously used to generate software entities from ontologies directly, they were limited to a few programming languages, were tightly coupled with specific implementation methods, and did not consistently support lightweight serialisation between data models and exchangeable documents. LLMs can provide cognitive solutions to real-world problems via similar model transformations. Especially for SBOL3, where the data model is complex with numerous entities and their relationships, such a hybrid approach can facilitate creating intuitive and user-friendly software libraries. Hence, integrating LLMs with ontologies can be valuable for large and graph-based models.

Here, we present an ontology-driven and LLM-based software engineering automation approach for knowledge graphs. In this software automation approach, a domain ontology acts as a metamodel to generate a corresponding software library, with the emphasis to make the library user-friendly. We initially present the SBOL3 ontology, which includes both specification-related and user-friendly terms. The ontology is further annotated according to the ontology-to-library (OtoL) framework, also presented here. We demonstrate this automation approach by creating the much-needed sbol-script library. This library supports server-side web development with TypeScript. It is also available as a distributable JavaScript file and can be run in a web browser. This approach facilitates the automation of software libraries in engineering biology, but it can also be applied to other graph-based data models for the sustainable development and maintenance of software projects.

## 2 Methods

This study presents an ontology-guided and prompt-oriented software generation methodology and applies it to create the sbol-script library. Our goal was not to train or fine tune an LLM but to evaluate whether semantically grounded ontology inputs could enhance iterative software generation for the graph-based SBOL standard. LLMs are well suited due to their capability to translate different types of data, including code, formal specifications in the form of ontologies, graph-based examples for validation, and iterative feedback to guide the creation of implementable software artefacts. This flexibility is important for this work, where the goal is not only to reproduce syntax but also to capture the necessary semantic relationships, constraints, and abstractions, and create maintainable developer interfaces. The selected LLM model was not treated as a standalone solution. Instead, it was incorporated as a tool into an ontology-driven formalisation workflow to enhance semantic reliability and reuse when creating software.

### 2.1 Generating ontologies

The SBOL3 and OtoL ontology definitions were developed using Owlready2 (*32*) in Python and were serialised into an RDF document. The ontology was then converted into the OWL and Manchester Syntax formats using the ROBOT library (*33*). An HTML version of the ontology was created with pyLODE (*34*). The cardinalities between entities are described via multiple approaches, including *functionality* of properties (0..1 or 1..1), *existential* restrictions (≥1) on core terms, and *domain* and *range* attributes of properties (0..*) (*23*). Cardinality restrictions were also used to additionally constrain properties that are used in multiple contexts.

### 2.2 Model selection and experimental workflow

Initially, both open-source and commercial LLMs were evaluated. Pilot experiments were conducted using different models (*35* –*37*) to observe improvements in context handling, code generation quality, iterative refinement of behaviour, semantic consistency, and time to execute a query. Open-source and self-hosted models, such as Llama and Qwen, were also considered. However, conducting these experiments with self-hosted models may require significant hardware resources, dedicated GPU infrastructure, inference engine support, memory optimisation, and continuous maintenance. Hence, for this work, commercial models (including Gemini 2.0, Claude Sonnet 4.5 and 4.6, and ChatGPT 4.5 and 5.4) were prioritised for the main experiment phase to focus on achievable implementations using the resources available.

### 2.3 Ontology-guided iterative generation workflow

The software generation workflow began with formalising domain requirements from SBOL3, using additional domain expertise. The requirements were captured in a machine-processable SBOL3 ontology. In addition to capturing the core entities and their relationships, the ontology was refined during the experiments by extending existing terms and adding new human-readable terms that are constrained using logical axioms and subclass relationships. These constraints form the basis of creating user-friendly software methods and classes.

The workflow methodology was implemented through a sequence of experiments, each representing a progressively more mature stage by extending the prompt used. Rather than treating each experiment as an isolated run, experiments were iteratively run to identify missing classes, incorrect abstractions, incomplete coverage, weak semantic descriptions, or usability issues in the generated output. These observations were then used to improve the subsequent prompt and ontology revision cycle.

### 2.4 Evaluation criteria

Generated outputs were evaluated throughout the iterative process using both semantic and practical criteria. These criteria included the implementation completeness of generated constructs, the semantic consistency with ontology defined entities and properties, support for higher-level abstractions, presence of required factory methods, execution behaviour of the generated code, usability of the resulting API, approximate response time, and token consumption. The generated outputs were also compared against the expected representations derived from the ontology and existing SBOL3 examples to ensure that the generated library preserves the intended syntax and semantic structure of SBOL3.

## 3 Results

We present an MDSE approach, in which ontologies formalise knowledge graphs for a domain of interest, and LLMs are then used to generate a corresponding software library (Figure 1). Our goal is not only to create the necessary methods to work with a defined data model, but also to facilitate the creation of developer-friendly APIs. Such APIs provide an abstraction layer between low-level entities that require domain expertise to use and high-level entities that are more intuitive for software developers. This abstraction layer is controlled via the rich expressivity of ontologies, ontological class definitions (e.g. equivalentTo), and inheritance relationships (e.g. subClassOf) to guide LLMs. This MDSE approach was demonstrated for SBOL (Figure 3). Here, we introduce the SBOL3 ontology, which acts as a semantic backbone to link the conceptual structure of the SBOL3 standard and the generative capabilities of LLMs for unambiguous interpretation of the specification and improving the consistency of the results.

**Figure 1:**
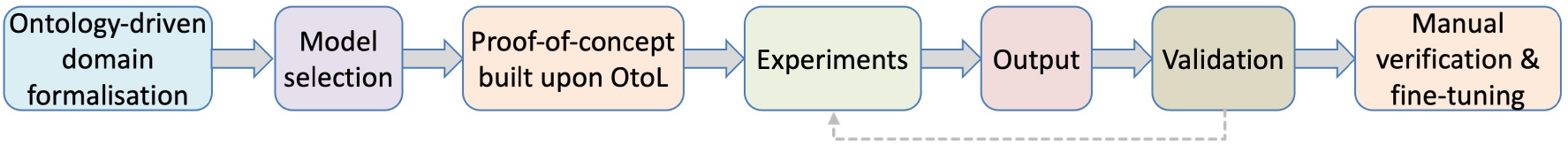
The ontology-driven software engineering approach using LLMs. The data model is initially captured as an ontology, which is then annotated and used as input to the selected LLM, together with a small proof-of-concept project based on a graph-based implementation pattern. An experiment and validation cycle is followed by manual optimisation and fine-tuning by a domain expert.

To formalise this MDSE approach, we present a lightweight framework, called ontology-to-language (OtoL), to control the implementation of software artefacts and annotate ontologies so that ontological definitions representing graph nodes and edges can be mapped to software entities and associated properties. Although OtoL was applied to the SBOL3 ontology to derive the sbol-script library, it can also be adapted for other software automation workflows.

Motivated by the increasing importance of web-based technologies and tools in the ecosystem of engineering biology, sbol-script can help tool developers who intend to create server- or client-side applications. Together, these contributions provide a practical outcome for the engineering biology community and a more general method for combining ontology engineering with LLM-assisted software development. Individual components of our workflow are explained in more detail in the following sections.

### 3.1 Ontology-to-Library framework

The ontology-to-library (OtoL) framework is a foundational component of the ontology-driven workflow. It includes a lightweight ontological layer to annotate domain-specific terms and an algorithm to represent graph-based entities within software artefacts. The key annotation properties include:

- `otol:domainEntity`: The annotated ontology term is represented as a first-class entity (e.g. classes) in the resulting library.
- `otol:constantList`: A defined ontology class, in relation to other terms, is annotated to generate a vocabulary list that is easy to access and use (e.g. software constructs with human-readable labels).
- `otol:replacementOf`: Controls how local terms inheriting from external terms are handled. The value of this property determines the name of the resulting software entity, while the annotated term is used to create it.

The OtoL framework controls the creation of software entities as wrappers that interact with an RDF knowledge graph without dealing with in-memory objects. This approach enables consistently compiling numerous ontological terms into software entities and constructs using ontologies and LLMs. The framework decouples the serialisation of knowledge graphs. This feature is particularly useful, as SBOL3 requires four graph serialisation types. The reference implementation has been developed in TypeScript and can be translated into other programming languages. The framework defines a base document class that stores a handle to the underlying knowledge graph. It also defines a base class for representing domain-specific entities, tracking the identifiers of corresponding graph nodes. For flexibility, generic setter and getter methods are included to set and extract values interacting with the corresponding graph nodes. The framework also includes methods to annotate nodes with custom properties and values.

### 3.2 SBOL3 ontology

The SBOL3 ontology has been designed as a machine-accessible representation of the specification, with over 200 SBOL terms to describe the necessary syntax and semantics. It significantly differs from the SBOL2 ontology (*23*) due to the major SBOL3 changes (*7*). The SBOL3 ontology incorporates over 100 additional terms from different ontologies, primarily to express design-related constraints, enrich designs, and align information with existing semantics. These ontologies include Sequence Ontology (SO), Systems Biology Ontology (SBO), the ontology of data analysis and management (EDAM), the Chemical Entities of Biological Interest (CHEBI) ontology and Gene Ontology (GO) (*7*). The SBOL3 ontology builds on a subset of the Provenance Ontology (PROV-O) and the Ontology of units of Measure (OM), mirroring the data model (*38, 39*). However, it decouples the imported entities by defining wrapper terms with appropriate inheritance and incorporating SBOL-specific information where necessary, without modifying external definitions. The resulting ontology is available in RDF and OWL formats, and can also be browsed through an HTML version.

The ontology defines terms for core SBOL entities (>40), relationships (>60), and properties (>20). The list includes top-level entities, such as `Component` and `Sequence`, which are information exchange containers, together with their child entities to capture contextual information (e.g. `SubComponent` and `Interaction`). The ontology introduces new terms when necessary, such as `GenericTopLevel` and `Metadata` for application-specific entities.

The ontology explicitly defines additional SBOL-specific terms (*∼* 45) to constrain and describe designs (e.g. `inline` or `reverseComplement` terms for orientation values). This information is often available in tables or as free-text in the specification. To reflect the specification and utilise the ontology for the MDSE approach, a parent term was also created for each related set (e.g. `Orientation`, `ConstraintRestriction`, and `Cardinality`). Similarly, parent terms (*∼* 13) were defined for related external ontological terms (e.g. `ComponentType`, `Encoding`, `InteractionType`, and `ModelLanguage`). For example, the `ComponentType` term is defined as a union of allowed values, such as `SBO:0000251` (DNA) and `SBO:0000252` (Protein). These *defined* terms form the basis for creating software constructs that can be accessed using labels rather than numeric identifiers, which can be difficult to remember.

The ontology provides additional helper terms (>35) that integrate information for specific contexts via multiple property-value assignments and inheritance layers (Figure 2). For example, `DNAComponent` derives from `Component` and has the predefined `SBO:0000251` (DNA) `type` value; `PromoterDNAComponent` derives from `DNAComponent` and sets the `role` property to `SO:0000167` (promoter); and `InChISequence` derives from `Sequence` and specifies the `encoding` property as `edam:format 1197` (International Chemical Identifier). Helper terms can include additional layers of inheritance. For example, `NonCovalentBindingInteraction` defines the interaction type as `SBO:0000177` and includes existential restrictions to `Reactant Participation` and `ProductParticipation`, which are further defined semantically. These helper terms are utilised to enhance the user experience.

**Figure 2:**
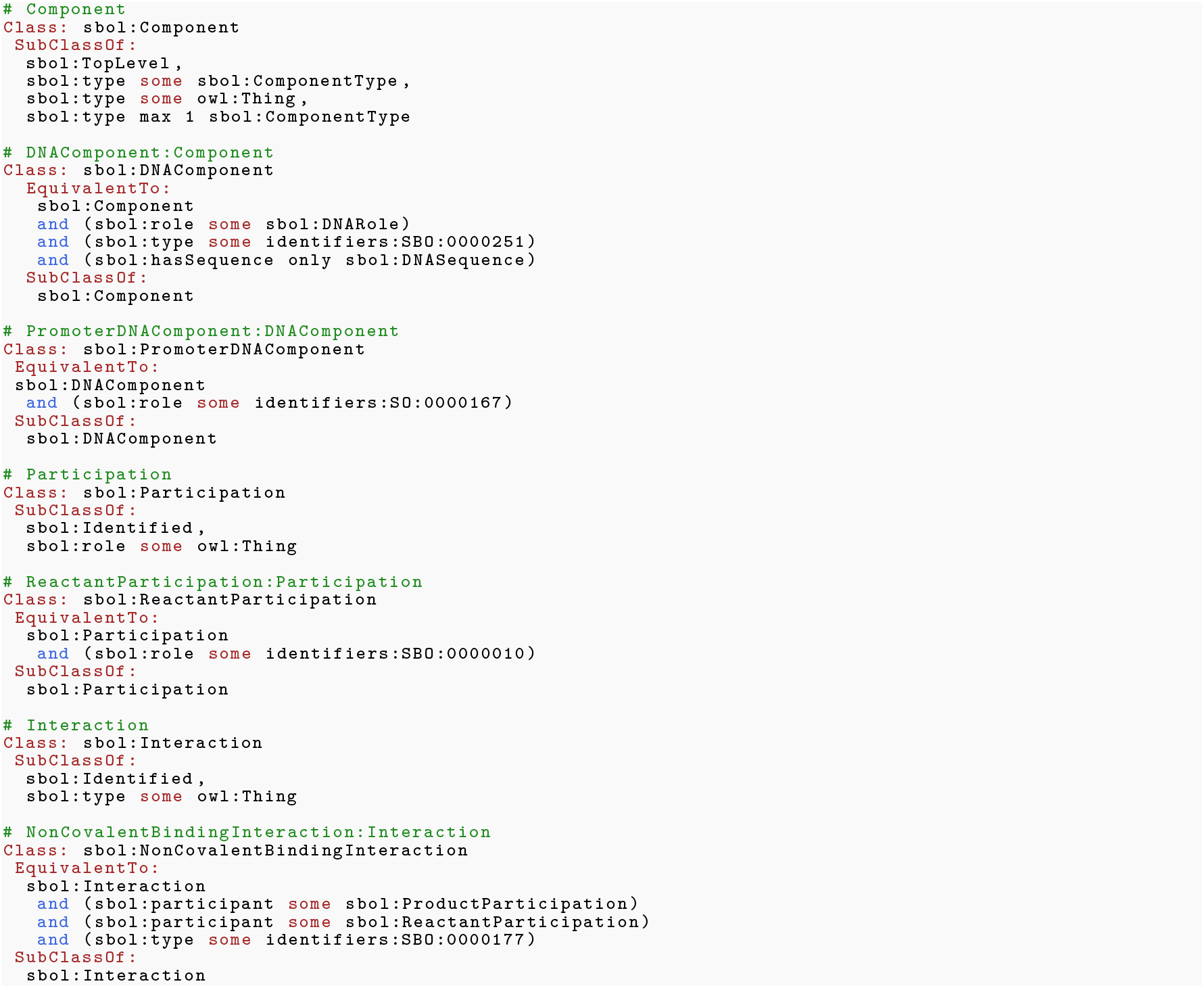
Examples of SBOL3 terms using the OWL Manchester syntax. The figure shows how inheritance is used to include additional information at different levels.

### 3.3 Iterative generation process

The software automation process relies on formalising the knowledge graph specification and a well-defined software pattern to generate the desired software artefact. In this context, the SBOL3 ontology formalises the SBOL standard and related requirements and provides a higher-level abstraction. The resulting semantic layer, the OtoL framework implementation in TypeScript, and the proof-of-concept demonstrating the implementation pattern using OtoL (for a few entities and properties only) were used as structured input to the selected LLM. The LLM was then prompted to scale the proof-of-concept into a complete software library, preserving the intended serialisation behaviour and ensuring that the generated classes accurately represent the domain entities, object properties, and abstractions defined in the ontology. A new TypeScript file was created for each core SBOL entity, and the existing files were populated accordingly.

Among the commercial models, Claude Sonnet 4.6 was selected as the most suitable model. The decision was based on comparative pilot experiments considering specific evaluation criteria (see the evaluation criteria in methods). It produced more stable outputs when aligning the ontology-defined requirements and the predefined implementation structure. Its performance showed the best overall balance for semantic consistency, stability during iterative refinement, implementation completeness, and practical execution efficiency.

The workflow was implemented iteratively (Figure 3). After each experiment, the generated outputs were compared against expected SBOL3 structures and behaviour, as well as representative SBOL3 examples. Revisions were then introduced through further prompting, with particular attention to the evaluation criteria (see Methods). Deficiencies identified in one iteration were translated into revised instructions for the next one. For example, in a later experiment, the prompt was extended to explicitly require adding factory methods, which are intended for creating child entities, since the previous iteration had failed to generate such methods. Hence, the prompt evolved from a general code-generation instruction set into a semantically constrained and implementation-oriented protocol.

**Figure 3:**
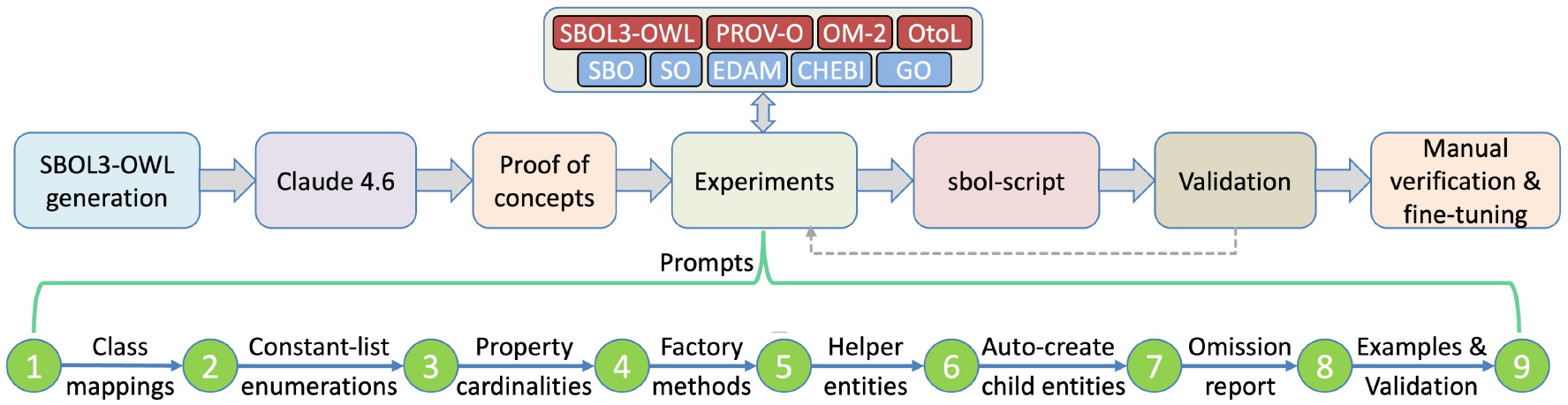
The workflow starts with identifying components of the proposed workflow. In total, nine experiments were used to iteratively generate sbol-script, followed up by a manual fine-tuning process.

During the mature experimental stage, the prompts were revised to incorporate specific requirements, such as: (i) ensuring to create TypeScript files for ontology classes annotated as domain entities, (ii) generating code for higher-level ontology classes, (iii) deriving controlled vocabulary constructs from ontology classes annotated as constant lists, (iv) generating factory methods for child entities based on object properties and selected restriction patterns, and (v) creating required subordinate entities where higher-level constructs implied their existence. In addition, the workflow required the model to report any omitted classes and properties, together with the reasons for exclusion, in order to identify any implementation gaps.

The experiments also included tasks to test whether the generated library could be used in practice by generating targeted examples. These tasks helped identify any limitations that were not always visible from static code inspection alone. This maturity-driven methodology helps evolve the ontology, the prompts, and the implementation together.

### 3.4 The sbol-script library

The sbol-script library enables creating genetic circuit designs that can be exchanged between different software tools. It was specifically developed in TypeScript to support different types of software, including server- and client-side web applications and standalone tools. It can be integrated with popular web frameworks due to its native web support.

The library stores designs in an in-memory knowledge graph, building on the OtoL framework. Properties of entities are accessed via simple getters and setters that interact with the underlying graph. The library can import and export designs using SBOL3 and supports all required file formats, including RDF/XML, Turtle, N-Triples and JSON-LD. The N-Triples format is particularly useful for streaming or comparing designs, as the library can also create ordered N-Triples.

Sbol-script manages designs via `SBOLDocument`, which can instantiate other entities as required. Class inheritance, cardinalities, and properties are derived from the SBOL3 ontology. Base classes, such as `Identified`, represent common behaviour. For example, `Identified` provides common properties such as name, description, source, related processes, and measurements. Moreover, user-friendly classes (e.g. `NonCovalentBindingInteraction`, `PromoterComponent`, and `StimulatorParticipation`) offer readily available functionality. Entity and property names and enumeration constructs are controlled via the `Vocabulary` class. For example, a DNA component’s type can be set to ‘`ComponentType.DNA`’ and serialised as an Internationalized Resource Identifier (IRI). Custom IRIs can also be used. In addition, the library includes an API layer, translated from the Java implementation, to create multiple parent and child entities and properties using utility methods that abstract complex operations. For example, the method to append a part to a device creates the required parent-child relationship and extends the device sequence.

### 3.5 Evaluation

Existing SBOL3 examples were used to evaluate sbol-script. These examples, including six use cases and 25 other examples, were previously created using libSBOLj3 (*11*), and were also used for pySBOL3 (*12*) development. Initially, a unit test was created for each example using getters and setters, considering a server-side implementation. Turtle, RDF/XML, N-Triples, ordered N-Triples, and JSON-LD files were created for each example. At the end of all experiments, when the resulting and original ordered N-Triples files were compared, all files were identical, except one, which included the creation of a random Uniform Resource Name (URN) as an entity identifier. To demonstrate the client-side use of the library, a local HTML page was created for each example with text areas for five serialisation outputs. The higher-level API was additionally tested using six use cases, and the resulting files were identical. These tests demonstrate the capability of sbol-script for different development goals.

## 4 Discussion

This work presents an ontology-driven approach that utilises LLMs to generate software libraries for knowledge graphs. The practical outcome of this work is the creation of reusable resources for the engineering biology community. First, we presented the SBOL3 ontology as a metamodel to capture the semantics and guide the software generation process. We then presented the OtoL framework to identify ontological terms that become software entities and control the creation of these entities. Finally, we demonstrated the approach to generate the sbol-script library, which can be incorporated into various applications.

In this approach, an LLM acts as the generator and a domain ontology serves as a metamodel that constrains the output. Once the ontology is annotated and the LLM prompt is reasonably stable, choosing the target programming language becomes a downstream concern. The same ontology and prompt can be used to create libraries in different languages, analogous to compiling bytecode for different platforms. Hence, regenerating libraries from a single semantic source removes recurring duplication of effort and facilitates sustainable software engineering.

Software sustainability is an important aspect of this work. Research communities often stay behind due to a lack of dedicated funding and maintenance support. Traditional software automation can be more controlled, typically involving tightly-coupled transformation definitions via manually maintained templates. However, such an automation is difficult to sustain once the specification begins to change. The LLM approach we present here offers flexibility. It can handle mixed inputs and produce implementable artefacts with significantly less manual effort. Moreover, the ontology-driven workflow shifts the maintenance effort: the ontology is updated, and the library is regenerated. This approach also scales predictably. As a result, what started as an *ad hoc* assistant became a reliable generator, and the ontology was rich enough to guide the end-to-end library generation, evolving a small proof-of-concept to cover the full SBOL3 data model. This methodology can be applied to any graph-based standard whose semantics can be represented in an ontology.

The methodology matured over iterations. Initially, the ontology and prompts could only support a few representative examples. To our surprise, the earlier experiments generated all 31 examples identically, only to realise that the LLM had hardcoded the file contents. Hence, all outputs were carefully investigated after each experiment. By the final iteration, all 31 reference examples could be regenerated almost identically, although there were small issues that required manual fine-tuning to create identical examples. Some of the issues were due to the LLM’s behaviour. For example, the LLM used a different capitalisation for the EDAM ontology, although there were no similar issues with other imported ontologies. Moreover, we noticed that the ontology construction process introduced some errors that propagated into the implementation. For example, the definition of `ControlInteraction` term had to be updated.

For knowledge graphs, incorporating ontologies in an LLM-based workflow is essential. SBOL3 includes definitions for several entities, relationships and cardinality rules. We initially tried to incorporate these requirements into the prompts. However, prompts grew past sensible limits, relationships silently disappeared, and minor wording changes cascaded into structural issues. Utilising the SBOL3 ontology in the workflow provided a stable and machine-readable foundation, and the OtoL annotations provided a complementary layer to specify mappings between the SBOL3 terms and software entities. An experimental prompt then acts as a contract over the ontology, not a restatement of a knowledge graph specification.

A side effect of utilising an LLM is not being able to anticipate how developer-friendly the resulting library will be. Here, we provided an ontological solution to this challenge by creating defined classes as helper terms. These domain-specific terms are integrated into the library to enhance the user experience. For example, terms such as `PromoterDNAComponent` or `NonCovalentBindingInteraction` are defined, wired with required properties, such as types, roles, and child entities. The `Vocabulary` class, derived from the constant-list annotations, replaces IRIs with human-readable enumerations. There are also additional challenges. The quality of the resulting library depends on the underlying ontology. Moreover, missing or ambiguous terms affect the implementation. Additionally, manual verification to assess the results is still required.

This work closes the gap between evolving community standards and subsequent software development. Our workflow can be transferred to any graph-based data model with an associated ontology. Hence, with a carefully crafted and annotated ontology, a stable prompt contract, and an iterative refinement process, complex graph-based data models can be turned into developer-friendly libraries. The SBOL3 ontology and the sbol-script library are also direct contributions to the community. In addition, we expect that the OtoL framework and the iterative refinement protocol defined here will be valuable in domains with similar interoperability and sustainability challenges.

## Data availability

The data underlying this article are available on GitHub at https://github.com/SynBioDex/sbol-owl3 and https://github.com/SynBioDex/sbol-script open-source repositories.

## Author contributions statement

All authors contributed to the manuscript. G.M. initially developed the ontology with contributions from the co-authors, developed the proof-of-concept framework, and optimised the final sbol-script library. M.U., T.M. and G.M. extended the SBOL ontology to be used by LLMs. T.M. and G.M. designed and carried out the LLM experiments.

## Acknowledgement

We thank the SBOL community. For the purposes of open access, the author has applied a Creative Commons Attribution (CC-BY) license to any Accepted Author Manuscript version arising from this submission.

